# Distribution of nematocytes differs in two types of gonophores in hydrozoan *Sarsia lovenii*

**DOI:** 10.1101/2023.03.22.533798

**Authors:** Alexandra A. Vetrova, Andrey A. Prudkovsky, Stanislav V. Kremnyov

## Abstract

Hydrozoan cnidarians are widely known for a diversity of life cycles. While some hydrozoan polyps produce medusae, in most species the gonophore remains attached to the polyp. Little is known about the mechanisms behind the loss of the medusal stage in hydrozoans. Hydrozoan *Sarsia lovenii* is a promising model for studying this issue. It is a polymorphic species with several haplogroups. One haplogroup produces attached eumedusoids and the other one buds free-swimming medusae. Here, we compared patterns of cell proliferation and distribution of nematocytes in medusoids, medusa buds and medusae of *S. lovenii*.

Cell proliferation is absent from exumbrella of late medusa buds and medusae, but presumably i-cells proliferate in exumbrella of medusoids. In exumbrella of medusoids, we also observed evenly distributed nematocytes with capsules and expression of late nematogenesis-associated gene, *Nowa*. Nematocyte capsules and *Nowa* expression were also observed in exumbrella of medusa bud, but we did not detect prominent *Nowa* signal in the bell of developed medusa. It is also known that abundance of exumbrellar nematocysts signs immaturity in medusae of *Sarsia* genus. Our data demonstrate that nematocyte distribution and associated gene expression in medusoids resemble medusa buds rather than developed medusae. Thus, sexually mature medusoids exhibit juvenile somatic characters, demonstrating signs of neoteny.

**Research highlights:** Hydrozoan *Sarsia lovenii* has attached eumedusoids and free-swimming medusae. The distribution of nematocytes in eumedusoids resembles that in medusa buds. This may indicate neoteny of eumedusoids.

## Introduction

Hydrozoa is a cnidarian taxon known for complex and diverse life cycles. In many hydrozoans, the life cycle includes benthic polyp stage which asexually produce sexual zooids (gonophores) by means of lateral budding. In some species, gonophore develops into medusa that detaches from the polyp and swims freely in the water column, where it feeds and grows until sexual maturity (Cartwright & Nawrocki, 2010). The presence of a medusa stage has been asserted to be an ancestral state for Hydrozoa (Marques & Collins, 2004; Kayal et al., 2018). However, the evolutionary tendency towards the reduction of the medusa stage is widespread among hydrozoans. In most hydrozoans, gonophores lack several important morphological features of medusae, and they spawn gametes, being attached to the polyp. Depending on the species, they exhibit varying degrees of morphological reduction. It ranges from medusoids that may have some medusa-like features such as remnants of tentacles, canal system, and gut/mouth, to sporosacs that are completely devoid of medusa structures (Cartwright & Nawrocki, 2010).

Though reduced gonophores independently evolved at least 70 times in hydrozoan lineage (Miglietta & Cunningham, 2012), our knowledge on mechanisms of the medusae reduction phenomenon is limited. It derives mainly from the phylogenetic analysis (Leclère, Schuchert, Cruaud, Couloux, & Manuel, 2009; Miglietta & Cunningham, 2012) and studies on gene expression in several species (Sanders & Cartwright, 2015a; Sanders & Cartwright, 2015b; Leclère et al., 2019). But investigations are mostly focused on fully developed medusae (e.g., Müller, Yanze, Schmid, & Spring, 1999; Spring et al., 2000; Reber-Müller et al., 2006; Kraus, Fredman, Wang, Khalturin, & Technau, 2015) while reduced gonophores are understudied besides morphological descriptions (e.g., Bourmaud & Gravier-Bonnet, 2005). Still, recent studies on the latter suggest that Wnt pathway may play a key role in the loss of medusa stage (Sanders & Cartwright, 2015a). The comparative study of differential gene expression between hydrozoans which exhibit the “book-end” phenotypes of gonophore development (fully developed medusa in *Podocoryne carnea* and sporosac in *Hydractinia simbiolongicarpus*) also revealed many genes that may participate in reduction of medusa stage. However, there are nearly 100 million years of divergence between these two species (Sanders & Cartwright, 2015b). To gain a better understanding of developmental, functional, and morphological differences between fully developed and reduced gonophores, the comparison of closely related taxa could be informative.

A potential model system for examining the medusae reduction phenomenon was introduced in the recent study (Prudkovsky, Ekimova, & Neretina, 2019). Hydrozoan *Sarsia lovenii* is a polymorphic species on genetic and phenotypic levels. It exhibit two different types of gonophores: one haplogroup produces attached eumedusoids and another one produces free-swimming medusae (Figure 1a). Medusoids (Figure 1b-c) possess umbrella and canal system but they don’t have functional mouth, tentacles and ocelli (Figure 1d) (Edwards, 1978; Prudkovsky et al., 2019). Colonies of *S. lovenii* also produce medusa buds which develop into medusae. Medusa buds have developing tentacle bulbs and tentacles tucked into the umbrella (Figure 1e) until medusae are fully formed (Figure 1f-g).

**Figure 1.**
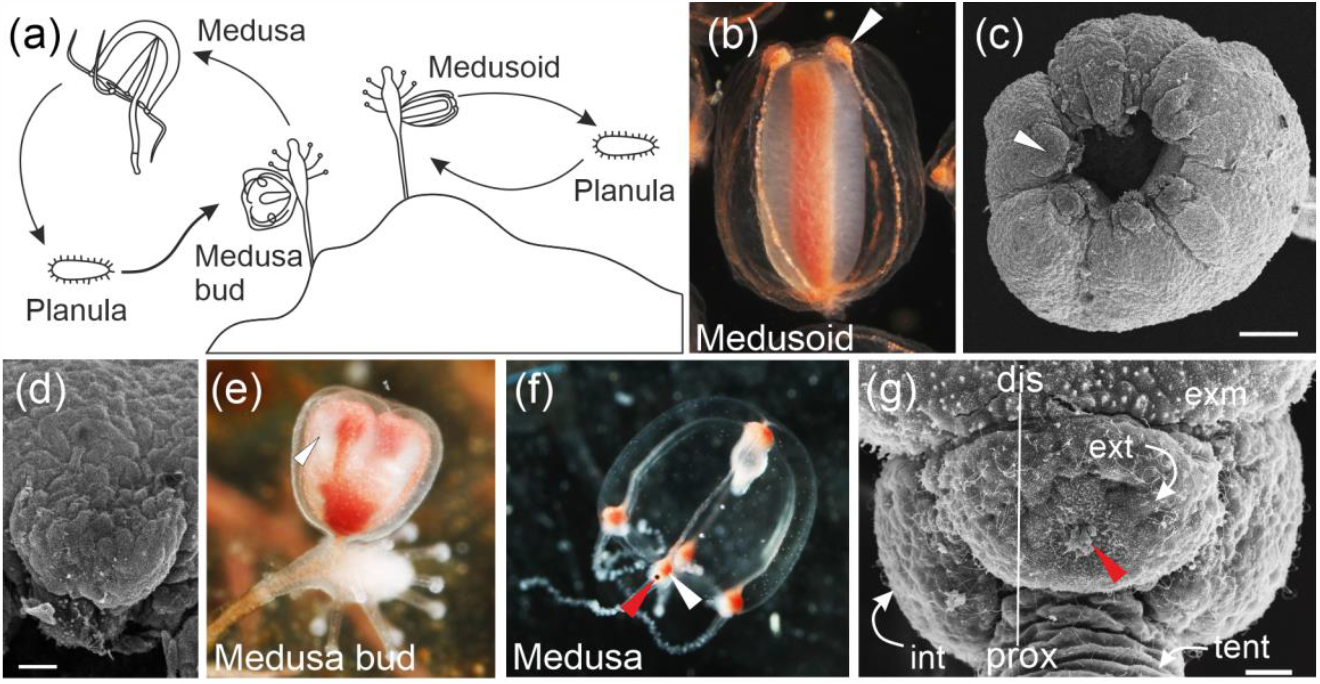
Gonophores of *S. lovenii*. (a) Life cycles of hydrozoans with free swimming medusa and reduced medusoid. (b) Medusoid. White arrowhead points to the tentacle bulb here and below. (c) SEM of the medusoid, oral pole view. (d) SEM of the medusoid’s tentacle bulb. (e) Medusa bud. (f) Young medusa. Red arrowhead points to the ocellus here and below. (g) SEM of the tentacle bulb. Dis – distal; prox – proximal; ext – external side of the bulb; int – internal side of the bulb; exm – exumbrella; tent – tentacle. Scale bars: (c) - 100 μm; (d, g) - 20 μm.

In this manuscript, we compared patterns of cell proliferation and localisation of nematogenic regions in medusa buds, medusae and medusoids of *S. lovenii*. Nematocytes are highly derived neuro-sensory cells generated from multipotent i-cells (David & Gierer, 1974; Miljkovic-Licina, Gauchat, & Galliot, 2004). Hydrozoans use nematocytes for feeding and protection (Kass-Simon & Scappaticci, 2002). Since medusoids have reduced morphology and simpler behavior compared to free-swimming medusae, we assumed that they might have differences in the distribution of nematocytes and localisation of nematogenic regions.

We used the EdU-labeling technique to determine the regions of cell proliferation. To localise nematogenic regions, we used light microscopy and performed in situ hybridization on marker genes of i-cells and nematocytes.

## Results

We determined the regions of cell proliferation in medusoids, medusa buds, and medusae of *S. lovenii* by EdU-labeling technique. In medusoids, EdU-positive nuclei were distributed unevenly along the oral-aboral axis and predominantly were found in the distal-most region, but absent in the proximal-most region. EdU-positive nuclei were detected in the umbrella and in reduced tentacle bulbs (Figure 2a). In the medusa bud, a few labeled nuclei could be observed only in the developing tentacle bulbs (Figure 2b).

**Figure 2.**
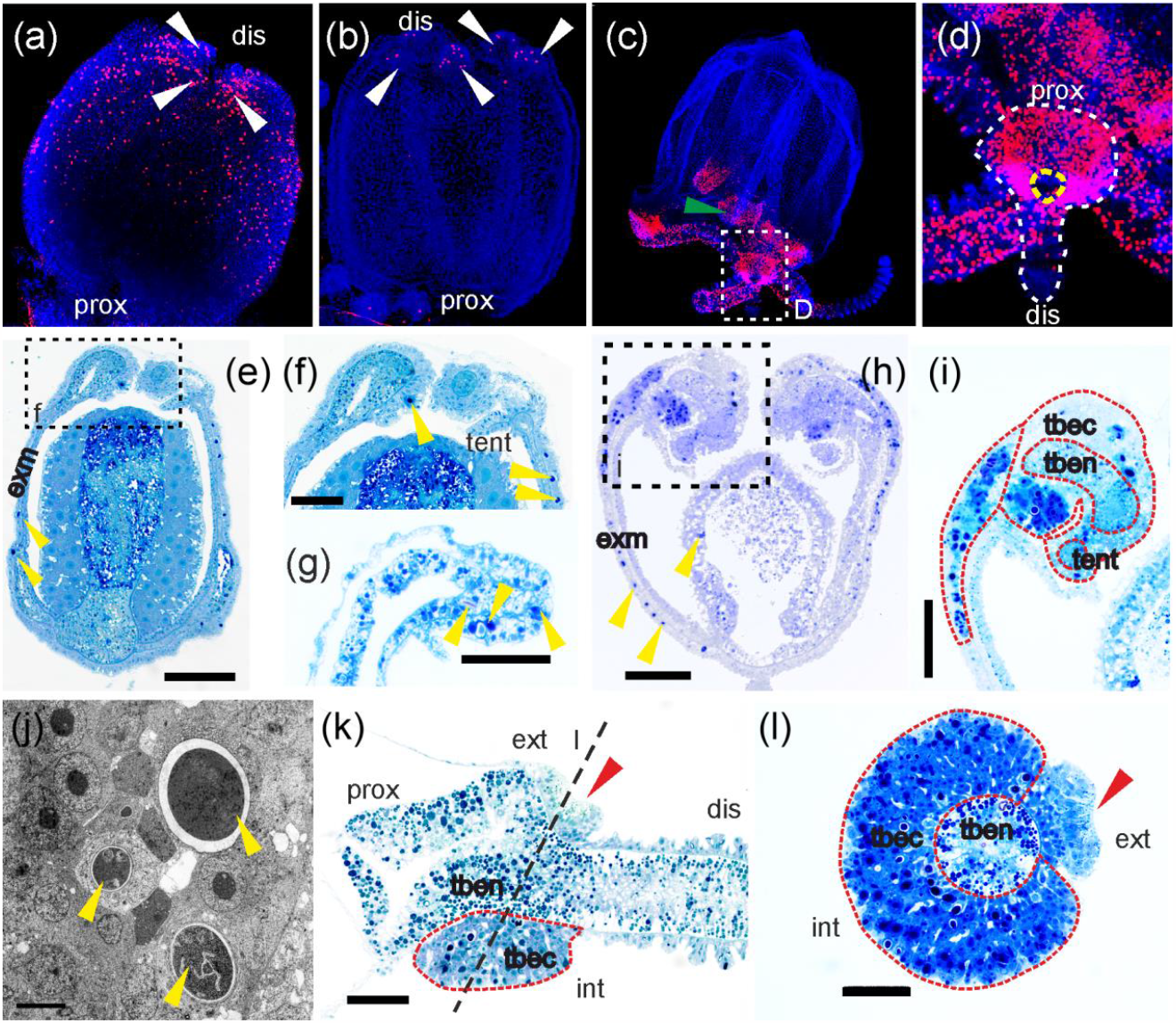
Cell proliferation (a-d) and nematogenesis (e-l) in gonophores of *S. lovenii*. (a) Medusoid. dis –distal, prox –proximal. (b) Medusa bud. (c) Young medusa. Green arrowhead indicate the proximal part of the manubrium. (d) White dotted line accentuate the shape of the tentacle bulb with a part of the tentacle. Yellow dotted line surronds the area free from EdU-positive nuclei (presumably, ocellus). (e, f) Semi-thin longitudinal section of the medusoid. Yellow arrowheads indicate nematocytes here and below. exm –exumbrella. (g) Semi-thin longitudinal section of the medusoid’s tentacle bulb. (h, i) Semi-thin longitudinal section of the medusa bud. Red dotted line accentuates nematogenic areas here and below. tbec – tentacle bulb ectoderm; tnen – tentacle bulb endoderm; tent – tentacle. (j) TEM of nematogenetic region. (k) Semi-thin longitudinal section through the tentacle bulb. Black dotted line shows the approximate section level on (l). ext –external side of the bulb; int - internal side of the bulb. (l) Semi-thin transverse section through the tentacle bulb. Scale bars: (e, h) – 100 μm; (f, k, l, i, g) – 50 μm; j – 10 μm.

In the medusae, intense EdU signal was detected in the very proximal part of the manubrium, in the proximal parts of tentacles, and in tentacle bulbs, except the area of the ocellus (Figure 2c-d). In the bulbs, EdU-signal intensity was lesser, proximal to the ocellus, suggesting that cell divisions are uneven (Figure 2d).

Further, we used semi-thin toluidine blue sections and TEM to examine the distribution of nematocytes in medusoids, medusa buds, and medusae of *S. loveni*. Toluidine blue allows easily distinguish nematocytes at various stages of differentiation from the other cell types by distinct blue coloration of the cytoplasm (Denker, Manuel, Leclère, Le Guyader, & Rabet, 2008).

In the medusoid, isolated nematocytes with capsules were found evenly distributed in the exumbrella (Figure 2e-f). Few nematocytes were also observed in the ecto- and the endoderm of a reduced tentacle bulb (Figure 2f-g).

In the medusa bud, nematocytes were found to be evenly distributed in the exumbrella except its distal-most part, adjacent to the developing tentacle bulb, where the region of active nematogenesis is clearly visible (Figure 2h-i). Another major nematogenic area is the ectoderm of the developing tentacle bulb itself. Transmission electron microscopy (TEM) demonstrated that nematogenesis stages ranging from multipotent i-cells (or early committed nematoblasts), characterized by a high nucleocytoplasmic ratio and a large nucleus with a prominent nucleolus (Chapman, 1974), to nematocytes with a full-sized capsule are present in this area (Figure 2j). Nematocytes were also found in the developing tentacle of the medusa bud (Figure 2j).

Generation of tentacular nematocytes continues in the tentacle bulb of the developed medusa. There, the most of the thickened ectoderm is nematogenic, except the area adjacent to the abaxial cup-shaped ocellus on the external side of the bulb (Figure 2k-l). Thus, nematogenic region in the tentacle bulb of *S. lovenii* is crescent-shaped as in *C. hemisphaerica* (Denker et al., 2008).

To determine the correspondence between regions of cell proliferation and regions of nematogenesis, we performed whole-mount *in situ* hybridization on medusoids, medusa buds, and medusae (Figure 3). Studies in *Hydra, C. hemisphaerica* and other cnidarians established a number of marker genes which expression is associated with multipotent i-cells and different stages of nematogenesis. We analyzed expression patterns of i-cells marker genes, *SlPiwi, SlVasa*, and *SlNanos1* (Mochizuki, Sano, Kobayashi, Nishimiya-Fujisawa, & Fujisawa, 2000; Mochizuki, Nishimiya-Fujisawa, & Fujisawa, 2001; Seipel et al., 2004; Rebscher, Volk, Teo, & Plickert, 2008; Leclère et al., 2012), and expression patterns of genes, associated with capsules of nematocysts, *SlMcol1* and *SlNowa* (Kurz, Holstein, Petri, Engel, & David, 1991; Engel et al., 2002; Denker et al., 2008). It was shown in *C. hemisphaerica*, that *Piwi, Vasa*, and *Nanos1* are also markers of early committed nematoblasts, and *Mcol1* is an earlier marker of nematoblast differentiation than *Nowa* (Denker et al., 2008; Leclère et al., 2012).

**Figure 3.**
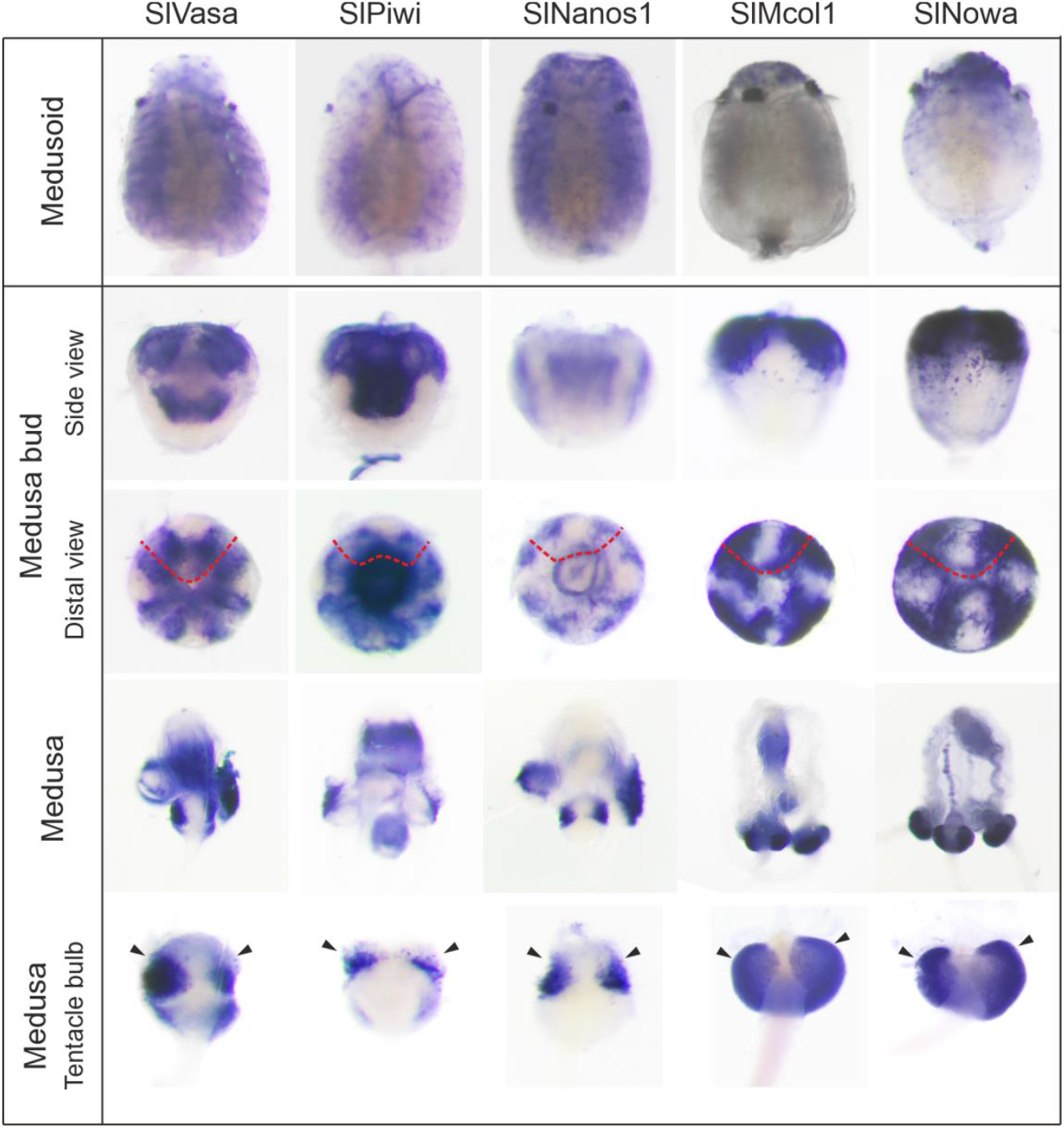
Spatial expression patterns of genes associated with i-cells proliferation (*SlVasa, SlPiwi*, and *SlNanos1*); genes associated with nematogenesis (SlMcol1, SlNowa) in medusoids (see the detail for an expression signal in reduced tentacle bulb), medusa buds, medusae and tentacle bulbs of medusae of *S. lovenii*. Red dotted lines outline wrapped tentacle bulbs. Black arrowheads indicate tentacle bulb insertion site on the umbrella. Medusoids and medusa buds directed with the distal end up. Bulbs are viewed from the external side. Note that manubria of medusoids protrude beyond the umbrella opening in case of *SlVasa, SlPiwi, SlMcol1*, and *SlNowa*.

In medusoids, scattered expression of *SlVasa, SlPiwi*, and *SlNanos1* was visualised in the umbrella except its proximal-most part where it attaches to a hydranth. Intense signal of all three genes was also detected in the manubrium ectoderm and in reduced tentacle bulbs (Figure 3). In medusa buds, their transcripts were detected in the manubrium area, likely, in the developing gonads. *SlVasa* and *SlPiwi* were also detected in the tentacle buds, wrapped inside the umbrella. However, the signal was absent from the central region on external side of the developing bulb. *Nanos1* signal was mostly observed in the umbrella, adjacent to tentacle bulbs (Figure 3).

In medusae, *SlVasa* and *SlPiwi*, but not *SlNanos1*, were detected in the manubrium area inside the umbrella, likely, in the gonad (Figure 3). In developed tentacle bulbs, *SlVasa, SlPiwi*, and *SlNanos1* expressing areas form a crescent around the bulb with a signal lacking from its external site (Figure 3). Equally intense *SlVasa* expression signal was detected in proximal and distal regions of the bulb but not in the most proximal region. *SlPiwi* exhibits intense signal in the proximal region of the bulb, similar to *SlVasa*, but the signal is less prominent in the distal region. *SlNanos1* transcripts were restricted to the proximal nematogenic ectoderm of tentacle bulbs (Figure 3).

Intense expression signal of structural nematocyte-accosiated genes, *SlMcol1* and *SlNowa*, was observed in the reduced tentacle bulbs and in the tip of the manubria in medusoids. *SlNowa* expression was also detected in scattered cells in the distal region of umbrella. In medusa buds, *SlMcol1* and *SlNowa* transcripts were detected in wrapped inside tentacle bulbs except the center of their dorsal side, in the adjacent umbrella regions, and in scattered cells in the umbrella (Figure 3). In medusae, *SlMcol1* and *SlNowa* was detected in the area of manubrium and in developed tentacle bulbs (Figure 3). In the latter, both genes were expressed through the entire nematogenic ectoderm in a crescent-shaped pattern (Figure 3).

## Discussion

In our study, we obtained patterns of cell proliferation, nematocyte distribution, and gene expression in medusoids, medusa buds and developed medusae of *S. lovenii*. Our results demonstrate that the distribution of nematocytes in medusoids resemble medusa buds. Numerous nematocytes (Figure 2e, h) as well as the expression of *SlNowa* (Figure 3), a marker of late nematoblast differentiation (Denker et al., 2008), were observed in the exumbrella of medusoids and medusa buds. Besides maturing nematoblasts, we also observed cell proliferation in the exumbrella of medusoids (Figure 2a). Since *SlPiwi, SlVasa*, and *SlNanos1* were also detected there in scattered cells (Figure 3), proliferating cells are likely i-cells/early committed nematoblasts (Denker et al., 2008; Leclère et al., 2012). However, in medusa buds and medusae, we didn’t observe cell proliferation (Figure 2b-c) and scattered expression of i-cells markers in exumbrella (Figure 3, S1). Likely, in late medusa buds, used in this study, early stages of exumbrellar nematogenesis are over already and are followed by later stages of nematoblast differentiation as *SlMcol1* and *SlNowa* expression indicate (Figure 3).

On the contrary, it is known that exumbrellar nematocytes are not abundant in medusae of *Sarsia* genus, being the characters of immaturity (Mayer, 1910; Edwards, 1978; Schuchert, 2001). We also didn’t detect expression of nematocyte marker genes and cell proliferation in the umbrella of developed medusae (Figure 2c, S1). Similarly, cell proliferation is absent from tissues of umbrella in late medusa buds and adult medusae in *Podocoryne carnea* (Spring et al., 2000). The possible interpretation is that medusoids exhibit not reduced, but juvenile characters on morphological (undeveloped tentacle bulbs, no mouth) and cellular (abundant exumbrellar nematocytes) levels. Thus, sexually mature medusoids of *S. lovenii* might fit with the definition of neoteny, supporting previously proposed explanation of medusa reduction in hydrozoans (Boero, Bouillon, Piraino, & Schmid, 1997). The presence of exumbrellar nematocytes may be an adaptive feature. Likely, they play a defensive role in attached medusoids, which are unable to move away from the threat.

We did not find localised regions of nematogenesis in medusoids (Figure 2e-g), likely, because they don’t have tentacles and mouth. However, we detected expression of all studied nematogenesis-associated genes in the area of reduced tentacle bulbs (tentacular nematocytes) and in the tip of manubrium (oral nematocytes) (Figure 3). The absence of *SlMcol1* expression, a marker of early nematoblast differentiation, from exumbrella (Figure 3) might indicate that progress of nematogenesis differs for exumbrellar, tentacular, and oral nematocytes in medusoids.

In medusa buds, we observed two regions of localised nematogenesis (Figure 2h-i), likely generating tentacular nematocytes. These regions differ in i-cells marker genes expression with *SlNanos1* mostly detected in the adjacent exumbrellar area and *SlVasa* and *SlPiwi* – in tentacle bulbs (Figure 3). In developed tentacle bulbs, we also observed differences in i-cells markers expression with *SlNanos1* absent from the distal region of the bulb (Figure 3). However, we didn’t detect such a difference in i.-cells markers expression in medusoids (Figure 3). Likely, there are several sub-populations of i-cells in *S. lovenii* The possible explanation is that the distribution of i-cells sub-populations is more ‘‘juvenile’’ in medusoids than in studied late medusa buds.

In tentacle bulbs of medusa buds and mature medusae, we detected expression of all examined nematogenesis markers (Figure 3). However, we didn’t observed ordered distribution of expression signal in contrast to the bulbs of *C. hemisphaerica*. In bulbs of *C. hemisphaerica*, i-cells marker genes are expressed in the proximal region, whereas cells expressing the *mcol1* gene are present in the intermediate area and *NOWA* is expressed in the distal part of the bulb (Denker et al., 2008; Leclère et al., 2012). Thus, the successive stages of tentacular nematocyte maturing are spatially ordered along a “cellular conveyor belt” in *C. hemisphaerica* (Denker et al., 2008). On the contrary in *S. lovenii, SlPiwi* and *SlVasa* are expressed in proximal and distal regions of the bulb, and *mcol1* and *NOWA* expression was detected in the whole nematogenic ectoderm (Figure 3). We also observed mixed up differentiation stages of nematoblasts in the developing tentacle bulb of a medusa bud (Figure 2j). Obtained results suggest that though the tentacle bulb is a conserved nematogenic site in hydrozoans, its organisation may vary in a species-specific manner. Nematogenetic site in the tentacle bulb of *S. lovenii* may be organised as in the *Hydra* polyp where differentiation stages of nematocytes are mixed up in the polyp body column, until maturing nematocytes migrate to their final destination (Bode, 1996). Further investigation will be needed to clarify this issue.

## Materials and Methods

### Animals and Sampling

Sampling of *S. lovenii* medusae and hydroids with gonophores at various stages of development was performed at the Pertsov White Sea Biological Station (Lomonosov Moscow State University) (Kandalaksha Bay; 66°340 N, 33°080 E). Medusae were sampled near the water surface close to the pier of WSBS and from the water column with a plankton net in adjacent localities in April-June. Hydroids were collected in the intertidal zone near WSBS. Medusa buds were found on the specimens collected in March-April, and medusoids were found on the specimens collected in May-June. Whole-mount observations were made under an Olympus SZ51 stereomicroscope. Whole-mount photos were taken with a Canon D550 camera supplied with a Canon macro 100 mm or Canon MP-E macro lens.

### Light and electron microscopy

For light and electron microscopy, specimens were fixed overnight at +4°C in a fixative solution containing 2.5% glutaraldehyde in the buffer solution (0.05 M cacodylate buffer with 15 mg/ml NaCl (pH=7.2–7.4)). Specimens were then postfixed in 1% osmium tetroxide in the buffer solution (2 hours for transmission electron microscopy (TEM), 15 min for scanning electron microscopy (SEM) at 22°C). They were then washed with the same buffer.

Further processing for histology and TEM was performed as previously described (Kupaeva, Vetrova, Kraus, & Kremnyov, 2018). Stained semi-thin sections were examined and photographed under Leica DM2500 (Leica, German) microscope equipped with a Leica DFC420C (5.0MP) digital camera. Contrasted ultrathin sections were examined with the JEM-1011 transmission electron microscope (JEOL, Japan). Proceedings for SEM were performed as described in Fritzenwanker, Genikhovich, Kraus, & Technau (2007). Some embryos were split into halves in 70% ethanol. Samples were examined under the JSM-6380 microscope (JEOL, Japan). Electron microscopy was performed at the Electron Microscopy Laboratory of the Biological Faculty of the Moscow State University.

### Examination of cell proliferation

For a visualization of cell proliferation, medusae, medusa buds, and medusoids were incubated in FSW (fresh sea water) with 20 μM EdU for 2 h at 14-16°C. Then, samples were washed 3x with filtered seawater and fixed in 4% paraformaldehyde in PBS overnight at 4°C. Further processing was performed according to the manufacturers’ protocol (Click-iT™ EdU Alexa Fluor™ 647 Imaging Kit, #C10340, Thermo Fisher Scientific). Samples were examined with a Nikon A1 confocal microscope (Tokyo, Japan). Z-projections were generated with NIS-Elements D4.50.00 software (Nikon).

### *S. lovenii* genes isolation, PCR, and antisense RNA probe synthesis

cDNA expression library was prepared by the SMART approach from total RNA with a Mint cDNA synthesis kit (Evrogen, Russia). cDNA gene fragments were isolated from the library by PCR with gene-specific primers. SlovMini1 dir: AATTACCGCTTCACTACTTTTCT, SlovMini1 rev: TACTGCCATCATATCATTTTCAC, SlovNOWA dir: TTTCTGGATGCTAATTGGCTACA, SlovNOWA rev: GGTGGTGGTGGTGGTGATG, SlovNanos1 dir: TGACGCATCGGTTTGTTTACTT, SlovNanos1 rev: TTCGATGCCATAGTAGCGTGT, SlovPiwi dir: TGTCTGGCCCCATTTTGTTG, SlovPiwi rev TGATGCGAAGTCTGTTGAGTGATT, SlovVasa dir: GAGCATTCGCCCAGTTGTAGTTTA, SlovVasa rev: ACCTCCGCCGCCATTCA

Primers were designed based upon sequences obtained from the sequenced transcriptome (Illumina) of *S. lovenii*. Transcriptome of *S. lovenii* was assembled and sequenced de novo and is available in our lab. Read quality control was performed with fastp (v.0.20.0) software (Chen, Zhou, Chen, & Gu 2018). De novo transcriptomes were assembled with rnaSPAdes (v.3.13.1) (Bankevich et al., 2012) software. Quality of assembly was assessed using BUSCO v.3.0.2 with metazoan database (Seppey, Manni, & Zdobnov, 2019).

Amplified fragments were cloned into the pAL-TA vector (Evrogen, Russia). Digoxygenine-labeled antisense RNA probes were generated from gene fragments, which were amplified from plasmids with *S. lovenii* genes.

### In situ hybridization

The *in situ* hybridization protocol was adapted from Vetrova et al. (2022). Samples were fixed with 4% paraformaldehyde in FSW overnight at +4°C, rinsed with PBS, and stored at -20°C in 100% methanol until hybridization. Samples were rehydrated with PTw (1x PBS with 0.1% Tween 20) and treated with proteinase K (80 μg/ml, 22°C) for 90 seconds. To inactivate the endogenous alkaline phosphatase and avoid a false positive result, samples were heated at +80°C for 30 minutes. Hybridization was performed at 58°C with digoxigenin-labelled antisense RNA probes (1ng/μL). Anti-DIG alkaline phosphatase-conjugated antibody (Roche; 1/2000 diluted) and NBT/BCIP substrate (Roche) were used to detect the probe. Stained samples were washed with PTw and methanol to reduce background staining and mounted in glycerol (87%). Imaging of samples was conducted using Leica M165C microscope (Leica, German) equipped with Leica DFC420C (5.0MP) digital camera.

### Image processing

Pictures were edited with ImageJ and Adobe Photoshop CS6 programs. Alterations to the “Brightness’’, “Contrast”, “Exposure”, and “Levels” for the RGB channel were used to achieve optimal exposure and contrast. All tools were applied to the entire image, not locally.

## Acknowledgments

We thank N.A. Pertsov White Sea Biological Station of Moscow State University for the help and support in obtaining samples and providing access to all required facilities and equipment of the “Center of Microscopy WSBS MSU”. We thank the Shared Facilities center “Electron Microscopy for Life Sciences” of the Lomonosov Moscow State University for the opportunity to perform electron microscopy. RFBR, 21-74-00129, supports the work.

## Conflict of interest

The authors declare no conflict of interest.

## Data availability

Sequences obtained in this study have been deposited in GenBank. SlovNOWA (OQ674077), SlovMini1 (OQ674078), SlovPIWI (OQ674079), SlovVASA (OQ674080), SlovNanos1 (OQ674081).

## Notes

### Competing Interest Statement

The authors have declared no competing interest.

